# Spontaneous activity changes in large-scale cortical networks in older adults couple to distinct hemodynamic morphology

**DOI:** 10.1101/2020.05.05.079749

**Authors:** Tatiana Sitnikova, Jeremy W. Hughes, Casey M. Howard, Kimberly A. Stephens, Mark W. Woolrich, David H. Salat

## Abstract

Neurovascular coupling is a dynamic core mechanism supporting brain energy demand. Therefore, even spontaneous changes in neural activity are expected to evoke a vascular hemodynamic response (HDR). Here, we developed a novel procedure for estimating transient states in intrinsic activity of neural networks based on source-localized electroencephalogram in combination with HDR estimation based on simultaneous rapid-acquisition functional magnetic resonance imaging. We demonstrate a readily apparent spatiotemporal correspondence between electrophysiological and HDR signals, describing for the first time how features of neurovascular coupling may differ among large-scale brain networks. In the default mode network, the HDR pattern in our older adult participants was associated with a surrogate marker of cerebrovascular deterioration and predicted alterations in temporal structure of fast intrinsic electrophysiological activity linked to memory decline. These results show the potential of our technique for making inferences about neural and vascular processes in higher-level cognitive networks in healthy and at-risk populations.

---

Intrinsic neural activity not explicitly associated with performing a task is an indicator of brain health. However, our understanding of how natural activity patterns arise is far from complete. The consequences of dysregulated intrinsic neuronal firing on life-sustaining cellular processes, such as gene expression and protein synthesis, are highlighted by animal models^1^. In the human brain, large-scale recordings at the cell population level link abnormal intrinsic neural activity to the progression of brain disorders, such as Alzheimer’s dementia^2–4^. Much of what we know about intrinsic neural activity in humans comes from functional magnetic resonance imaging (fMRI), which relies on neurovascular coupling to make inferences about neural activity^5^. Inherently, variations not only in neural activity but also in properties of the associated vascular support, which is the signal detected by fMRI, can influence patterns in the data, especially in the aging brain prone to cerebrovascular changes and patients with brain disorders accompanied by vascular comorbidities^6–8^. Therefore, new procedures for disambiguating between the neural and hemodynamic features in the human brain are needed.

Studies using fMRI have demonstrated that large-scale brain networks recruited by tasks are also spontaneously reactivated during resting state^9–12^. Two recent analytic approaches, focusing on electrophysiological correlates of neural activity that are temporally resolved on rapid scales of cognition^13^, suggested that such spontaneous reactivations may constitute transient states of network activity coupled to local vascular hemodynamic response (HDR). First, a novel mathematical tool based on the Hidden Markov Model (HMM) discerned recurring transient states of spatially coordinated amplitude changes (on average 100-ms-long) in resting magnetoencephalographic (MEG) data in both young and older adults^14–16^. This method demonstrated topographic correspondence between the estimated cortical loci of the unique rapid activity states and the large-scale networks previously observed with resting fMRI, raising a question of how the two imaging modalities are linked in real time. A second, complimentary technique showed temporal concurrence between the features of electrophysiological and HDR signals through simultaneously recording electroencephalographic (EEG) and fMRI resting data^17–19^. However, reliance on the assumed ‘canonical’ HDR shape to relate brief EEG states, when non-identical scalp-signal distributions suggested distinct neural sources, to hemodynamic fluctuations in brain networks lessened generalizability of this method. The canonical HDR shape has been estimated in sensory/motor primary cortices in young healthy participants^20–23^ and may not accurately describe the HDR in other large-scale networks or participant populations. The goal of the present study was to build and improve on these prior efforts by assembling a data-driven technique for estimating rapid electrophysiological network activity and the associated HDR, with an intent to advance both basic and clinical investigation of the intrinsic brain activity in relation to neurovascular coupling.

This study extends our prior MEG-based application of the HMM^14–16^ to the resting high-density EEG, source-localized to 38 regions in the entire cortex. The HDR patterns associated with different amounts of time spent in a given HMM state during timeframes in the simultaneously acquired fMRI data are estimated without *a priori* assumptions about the canonical HDR shape – by using finite impulse response (FIR) basis functions^23^ and shape-flexible gamma function^22^. This methodology establishes both spatial and temporal correspondence between the electrophysiological and hemodynamic activity in cortical networks in the resting brain, as well as delineates the morphology of HDR evoked by distinct electrophysiological network states. When we previously described electrophysiological network states in older adults based on HMM applied to a resting MEG, we observed spatial differentiation between coordinated activity states with amplitudes higher or lower than the session average^16^. Here, we describe the HDR evoked to meet the metabolic demand associated with opposite-polarity modulations in electrophysiological amplitude of similar states, inferred in older adults based on EEG. We also assess the relationship between HDR patterns and the temporal structure of electrophysiological activity states, in the context of variable cerebrovascular and cognitive decline across older adults. Given that the brain disproportionately consumes energy compared with other organs^24^ but maintains almost no energy reserve^25^, the HDR that promptly supplies nutrients for energy homeostasis^26^ may be closely linked to intrinsic neural activity, and any de-coupling may disrupt the intrinsic neural patterns. Although we examined several cortical networks, we focused on the default mode network (DMN), which is implicated in episodic memory and is vulnerable to age-associated changes^27^, in comparison to the dorsal attention network (DAN), which tends to remain functionally intact in early stages of neurodegenerative disorders, such as Alzheimer’s dementia^28,29^.

## Results

### Brief network states in source-localized EEG were discerned using HMM

Electrophysiological states lasting only a fraction of a second were detected in resting-state high-density 256-channel EEG of 15 older adults, which was source-localized to 38 brain regions using beamforming^14^ with spatial signal leakage reduction via orthogonalization^30^. This was the first application of the HMM that enabled to discern transient states of coordinated cortical activity based on EEG. Figure 1 illustrates the basic principle of HMM state inference on a sample segment of data. States inferred by HMM feature spatially coordinated changes in the amplitude of band-limited (4-30 Hz) electrophysiological activity that recurred in unique brain networks at distinct points in time. Comparable results were observed when HMMs with 10, 12, and 14 states were inferred based on the time-courses of activity amplitudes concatenated across participants. The resulting HMM states had clear spatial correspondence across participants, but each participant displayed their own state time-course, indicating when different states were most probable.

**Figure 1.**
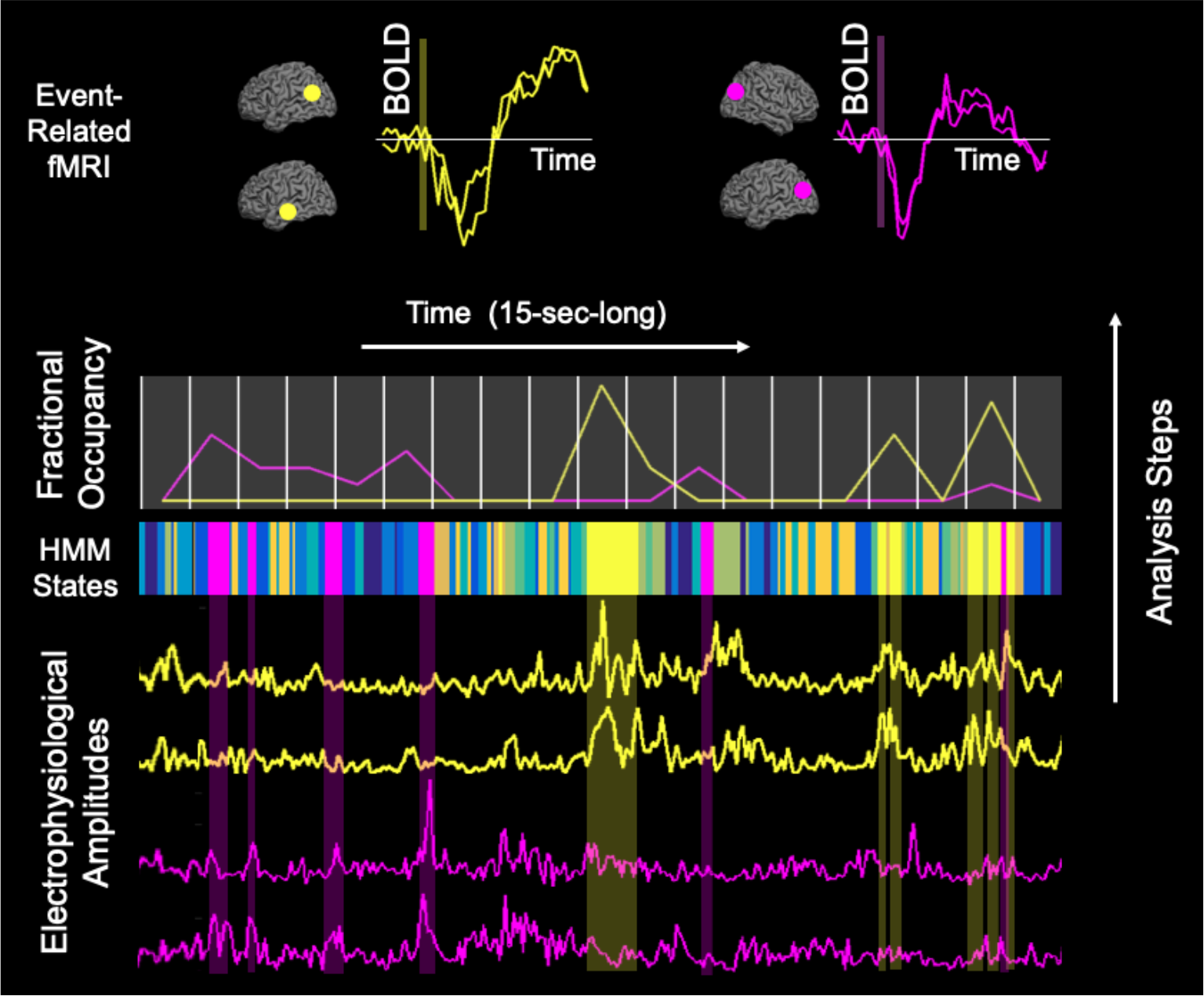
Analysis steps for simultaneously acquired EEG/fMRI data are illustrated based on a sample 15-s data segment from an individual participant (see Methods for further details). From bottom to top: HMM analysis applied to EEG-based electrophysiological amplitudes inferred a time-course of 10 recurring HMM states, which was converted into time-courses of fractional occupancy that were used as regressors to model neurally evoked HDR in event-related fMRI analysis. The multicolored band (middle panel) shows a sequence of inferred HMM states; 10 different colors correspond to 10 states characterized by distinct spatial patterns of electrophysiological activity. The yellow HMM state reflects rises in electrophysiological amplitudes (yellow time-courses in lower panel) that are source-localized to nodes of the DMN (yellow dots on cerebral cortex in upper panel). Likewise, the magenta HMM state reflects rises in (magenta) electrophysiological amplitudes that are source-localized to nodes of the visual network (VisN/magenta dots). Fractional occupancy was computed as the time in ms when each electrophysiological network state was active during each of 800-ms consecutive fMRI acquisition frames, demarcated by vertical white lines on a shaded background (middle panel); DMN/yellow and VisN/magenta time-courses of this fractional occupancy show different fluctuation patterns across frames. Deconvolved BOLD changes (left upper panel: DMN/yellow; right upper panel: VisN/magenta) were time-locked to ‘trigger’ frames with non-zero electrophysiological fractional occupancy, illustrated by vertical yellow/magenta shaded bars.

Cortical topographies of EEG-based HMM states resembled those previously observed by MEG in young^14^ and older adults^16^ using similar analysis methods and showed overlap with classic large-scale functional networks described based on coordinated hemodynamic fluctuations measured by resting-state fMRI^31–38^. State topographies were visualized by computing partial correlations between state time-courses and electrophysiological amplitudes in different cortical regions for each participant. Group maps were derived by averaging across participants. Figure 2 shows maps of five salient states (see Supplemental Figure 1 for maps of additional modeled states) indicating cortical regions that displayed higher or lower amplitudes in each state compared with average electrophysiological activity over time. State 1 was marked by increased amplitudes in regions overlapping the visual network (VisN), including the visual and posterior parietal cortices^14,33^. State 2 was marked by decreased amplitudes in regions overlapping the DAN, including the posterior lateral temporal cortex and intra-parietal sulcus and extending toward the frontal eye field^14,31,39,40^. State 3 was marked by increased amplitudes in regions overlapping the DMN, including the precuneus, posterior cingulate cortex, medial prefrontal cortex, temporal cortex, and inferior parietal lobe^32,35^. State 4 was marked by increased amplitudes in regions corresponding to the anterior DMN (aDMN), including medial and inferior-lateral prefrontal cortices and the temporal cortex^41^. Finally, state 5, which was clearly detected only in the 14-state HMM, was marked by amplitude increases in the sensorimotor network (SMN)^14,33,34^.

**Figure 2.**
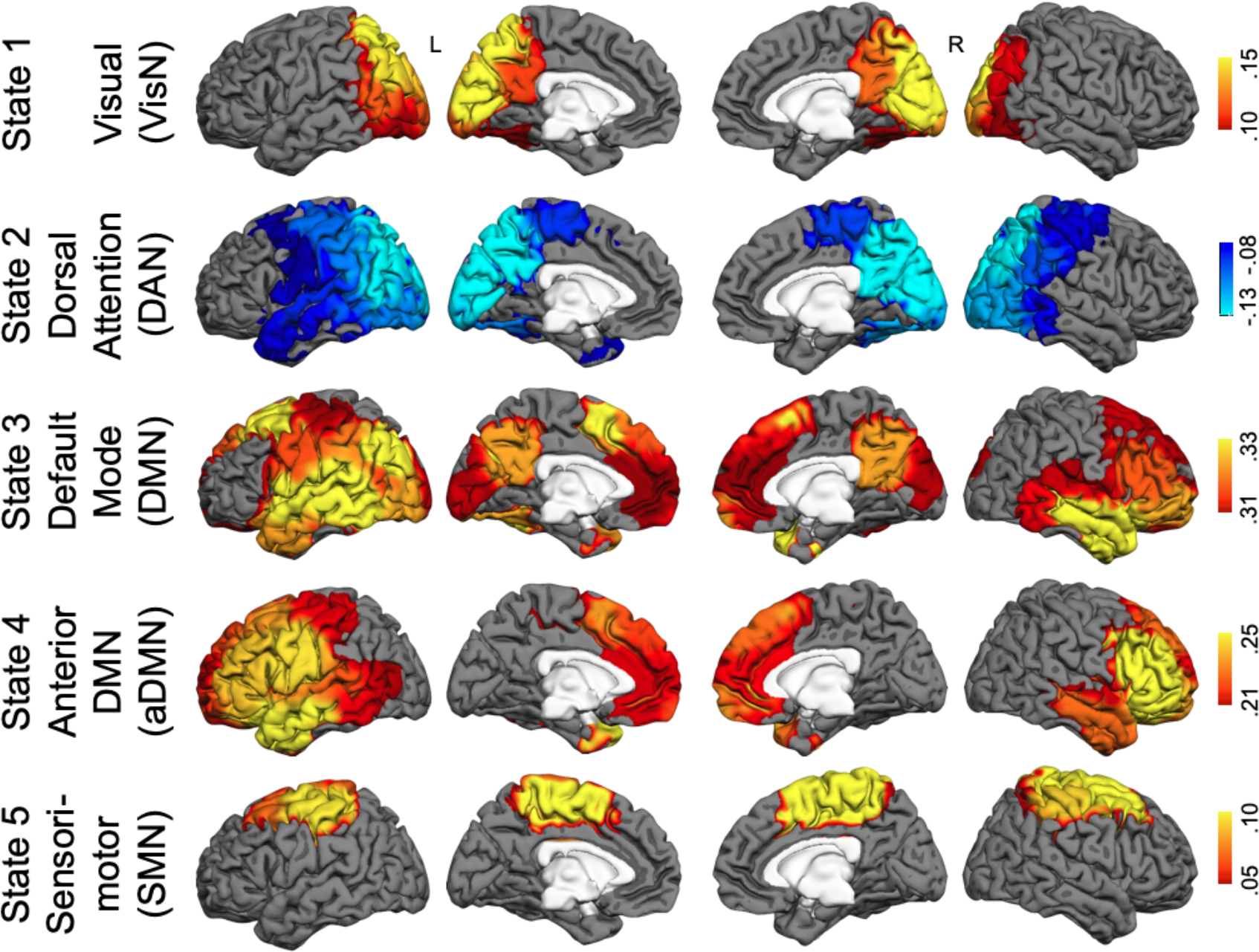
Maps of state-specific increases (yellow/red/brown) and decreases (blue) in electrophysiological amplitudes during five salient network states inferred by HMM. Maps show partial correlations between state timecourse and electrophysiological amplitudes in 38 ROIs combined across 15 subjects.

### Temporal pattern of electrophysiological activity states

Time-courses of EEG-based HMM states describe time-varying patterns of coordinated electrophysiological activity in unique neural networks at timescales under 100 ms. Figure 3 illustrates statistical properties of the durations of coordinated activity states in each neural network, here termed “lifetimes” (previously also referred to as “dwell times”^15,42,43^). Supplemental Figure 2 shows this information for the durations of intervals between the activity states in each network. In addition to mean and standard deviation, we computed the probability density function (PDF), which reflects likelihood of different lifetime/interval values during the recording period. Recent innovative analyses of electrophysiological recordings at micro and macro scales revealed temporal patterns in intrinsic neural activity with heavy-tailed PDFs, approaching the power law distribution^44–48^, which tend to spatially organize within commonly co-activated large-scale cortical modules^49^. These patterns may have physiological significance, as they are indicative of a system near the state of so-called criticality, in which available resources are kept at maximal availability for optimized responsiveness to the environment^50,51^. To describe the temporal activity structure for each HMM state-defined cortical network and for each participant, we used a statistic indicating the extent of deviation of the empirical PDFs from a fitted power law model^52^. When this statistic was computed on state lifetimes, a comparison of DMN/aDMN states to the DAN state, while regressing out the effects of age and sex, revealed differences in the extent of deviation from the power law (effect of state in omnibus repeated-measures analysis of variance [rmANOVA], F=21.90, p<0.05; follow-up: DMN<DAN, F=42.64, Cohen’s d=4.89, DMN<aDMN, F=9.64, d=0.68, aDMN<DAN, F=17.59, d=1.45, ps<0.05). Thus, the DMN state showed temporal activity structure closest to critical behavior. Interestingly, the DAN state characterized by reduced electrophysiological amplitudes, suggestive of reduced phase-locking of low-frequency activity^15^ and neuronal firing desynchronization^53^, showed the largest deviations from power law and presumed criticality, consistent with findings in cats and monkeys^53^. Similar results for intervals between the activity states in each network are reported in Supplemental Figure 2.

**Figure 3.**
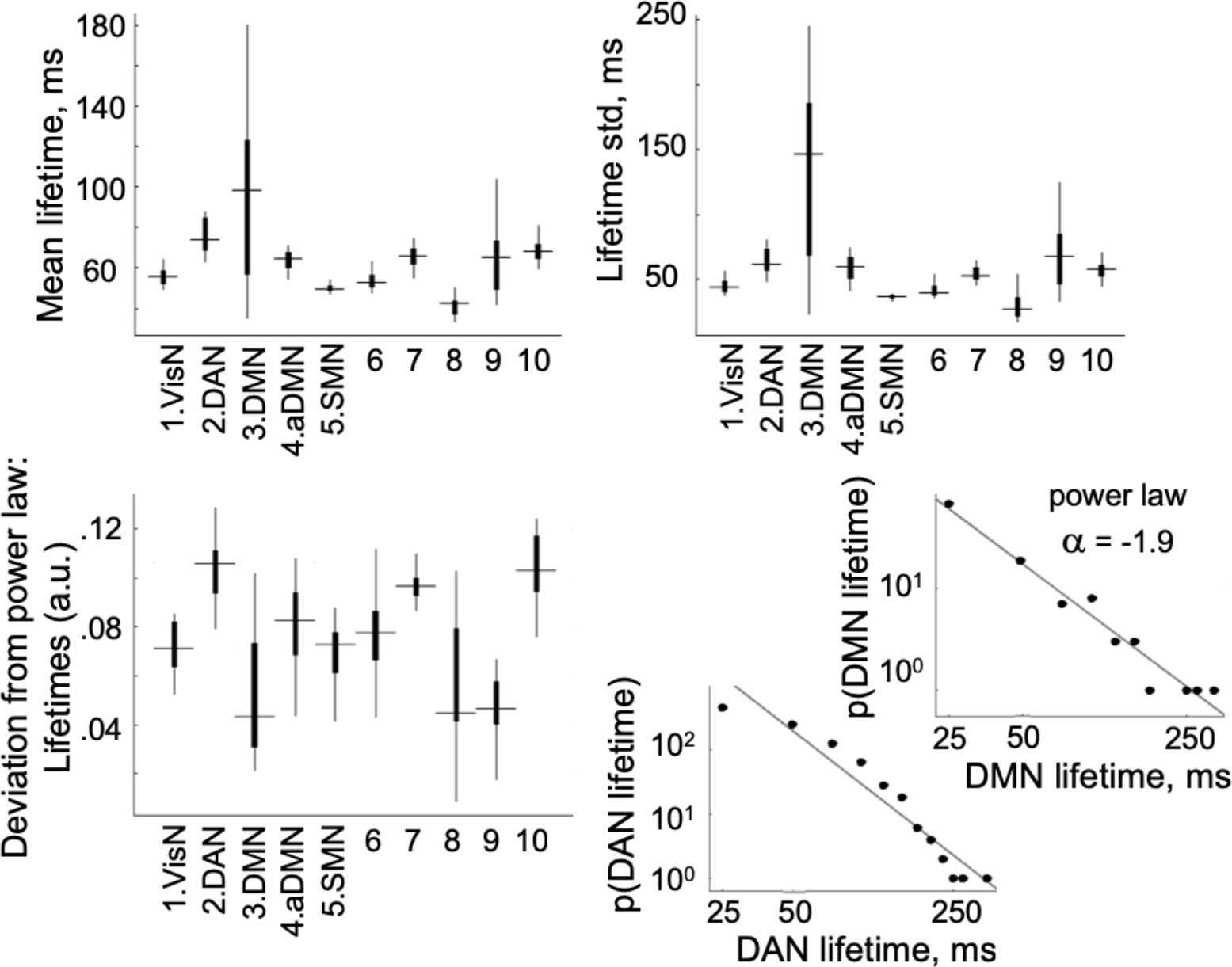
Temporal pattern of inferred HMM state lifetimes. For each state and participant, we computed the mean and standard deviation (std) of the coordinated activity lifetimes (upper panel). We also computed the deviation between empirical lifetime data and the power law function fitted using maximum likelihood estimation (lower left panel). Boxplots for the five salient states (1–5; shown in Figure 2) and five other modeled states (6–10; see Supplemental Figure 1 for maps of these states) summarize the group results (medians and 25th/75th percentiles; whiskers extend to the most extreme points not including outliers). Examples of lifetime distributions of DMN and DAN states from an individual participant (lower right panel). For DAN, the probability (p) of longer states was lower than would be expected for a power law distribution; for DMN, the grey line indicates the power law with exponent a=-1.9. Abbreviations of HMM state names are the same as in Figure 2.

### HDR evoked by time-varying fractional occupancy of electrophysiological network states

Because HMM electrophysiological states showed an irregular pattern of recurrence (as illustrated in Figure 1), the proportion of time spent in a given state during each 800-ms recording frame in simultaneously acquired fMRI data varied over time. This time-varying fractional occupancy per fMRI frame computed for each state was used as a predictor in general linear modeling (GLM) of state-specific changes in the blood oxygen level dependent (BOLD) signal recorded by fMRI, which is sensitive to regional neurovascular coupling^54^. HDR was quantified as the slope of BOLD change per unit increase in the electrophysiological fractional occupancy.

Figure 4 depicts maps of state-specific BOLD responses time-locked to fMRI frames with non-zero fractional occupancy by each of the five salient HMM states. The HDR, evoked by such electrophysiological “trigger-frames”, was modeled at each voxel using a gamma response function. The cortical maps show (de)activation clusters observed when group statistics were computed while treating participants as a random effect, regressing out the effects of age and gender, and correcting for multiple comparisons using simulation testing. The upper right panel summarizes shape parameters of the best-fit gamma models for each network node. Note that parameters for different nodes in the same network appear more similar to each other than those from different networks. After controlling for age/gender, the mean Euclidean pairwise distance between (*α, τ*) points, computed for each participant, is shorter within networks than between networks (F=16.86, p<0.05, Cohen’s d=1.31). This holds even though some of the nodes from different networks are proximal in the physical brain space (e.g., DMN state: inferior parietal, DAN state: intraparietal). The *α* exponent for all regions was >2 – the value commonly used in ‘canonical’ gamma response modeling^22,23^; noteworthy, this difference could be caused, in part, by slightly imprecise time-locking to electrophysiological states occurring at different times during the 800-ms trigger-frame. As expected based on the cortical topography of state-specific electrophysiological activity, the DAN state evoked a BOLD signal increase in bilateral inferior-parietal sulci and the posterior-temporal cortex (voxel-wise Cohen’s ds between 0.49 & 1.05). Similarly, regions where the BOLD signal was modulated in response to the DMN state corresponded to the electrophysiological topography of this state. Nonetheless, these regions, including the inferior-parietal lobe, precuneus/posterior cingulate cortex, medial/superior prefrontal cortex, and lateral-temporal cortex, showed an evoked BOLD decrease (ds between 0.51 & 0.97). This reversal in the sign of evoked BOLD changes corresponded to the reversal in the sign of electrophysiological amplitude changes—a decreased amplitude in the DAN state but increased amplitude in the DMN state (cf., Figure 2). The other states showing enhanced electrophysiological amplitudes also displayed spatiotemporally concordant reductions in BOLD signal. The VisN state was associated with BOLD reductions in regions of the dorsal visual stream^55^ (ds between 0.48 & 0.77). The aDMN state was associated with BOLD reductions in the medial, dorsal, and lateral prefrontal cortices as well as temporal and inferior parietal regions. The SMN state was associated with BOLD reductions in the sensorimotor cortex (ds between 0.43 & 0.79). Overall, the topography of cortical networks in which the BOLD signal was modulated by electrophysiological HMM states corresponded well to previous cortical parcellations into large-scale functional networks using variance decomposition techniques^56^ applied to continuous resting-state BOLD recordings^36^.

**Figure 4.**
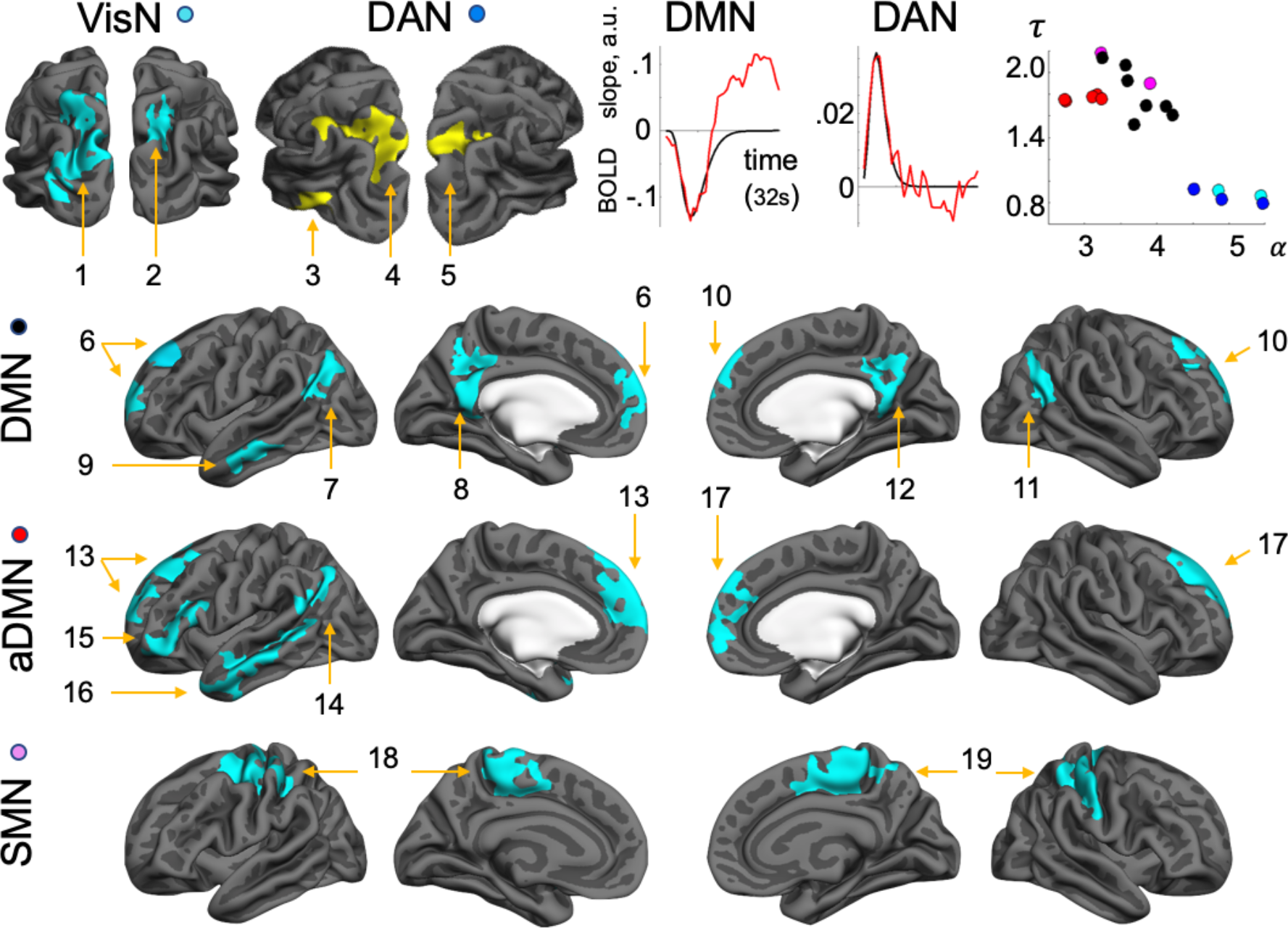
Maps of state-specific increases (yellow) and decreases (blue) in BOLD signal time-locked to ‘trigger’ timeframes with non-zero fractional occupancy by electrophysiological HMM states and modeled with a gamma response function. The slope of BOLD change in response to frame-to-frame fractional occupancy variation was estimated in each voxel. Upper right panel from right to left: shape parameters of the fitted gamma models (*α*, exponent, *τ*, dispersion) for VisN nodes in cyan, DAN in blue, DMN in black, aDMN in red, and SMN in magenta; graphs for DAN and DMN showing gamma models (black) with overlaid FIR-based BOLD signal estimates for representative nodes averaged across node voxels and participants (red). Numbers of significant clusters (p<0.05, corrected) represent: *for VisN state*, 1, left extrastriate and posterior parietal; 2, right posterior parietal; *for DAN state*, 3, left posterior temporal; 4 and 5, left and right intraparietal sulci and posterior parietal; *for DMN state*, 6 and 10, left and right medial/superior frontal; 7 and 11, left and right inferior-parietal; 8 and 12, left and right precuneus/posterior cingulate; 9, left lateral-temporal; *for aDMN state*, 13 and 17, left and right medial/superior frontal; 14, left inferior-parietal; 15, left lateral prefrontal; 16, left lateral temporal; *for SMN state*, 18 and 19, left and right sensorimotor and supplementary motor. HMM state name abbreviations are the same as in Figure 2.

Figure 5 shows time-courses of BOLD changes evoked by electrophysiological trigger-frames computed using a FIR approach^23^ without any *a priori* assumptions about HDR shape. For each network node, the values at each time-point reflect the average slope of BOLD signal modulation by changes, from trigger-frame to trigger-frame, in fractional occupancy of the eliciting electrophysiological HMM state. DAN, VisN, and SMN states evoked a monophasic response; after approximately 10 s, the BOLD signal returned to near-baseline levels. Noteworthy, for VisN and DAN states, a canonical gamma response function (*α*=2, *τ*=1.25) was well suited to model the HDR. We used a statistic indicating the extent of deviation of the observed BOLD time-courses from both fitted shape-flexible and canonical gamma response models. After controlling for age and gender, no differences between models were observed in all VisN and DAN nodes (Fs < 3.07, ps > 0.1, uncorrected). In the DMN, increased amplitudes of electrophysiological activity in lower frequencies (4-30 Hz) previously have been linked to higher BOLD levels^57,58^. Thus, the DMN state, characterized by increased lower-frequency amplitudes, would be expected to evoke a positive BOLD response. We observed a biphasic BOLD shape in several regions in response to the DMN/aDMN states; after an initial negative-going peak, the signal displayed a positive-going curve. In the second half of the epoch (16-32 s), the integral of the slope was significantly higher than baseline in a subset of DMN nodes (change from baseline: F=7.21, p<0.05, interaction of change from baseline with node: F=3.01, p<0.05; follow-up analysis revealed that six time-courses were higher than baseline; Cohen’s ds between 0.62 & 0.83). However, in a control analysis for DAN, VisN, and SMN, such effects were not significant (Fs<1, ps>0.35).

**Figure 5.**
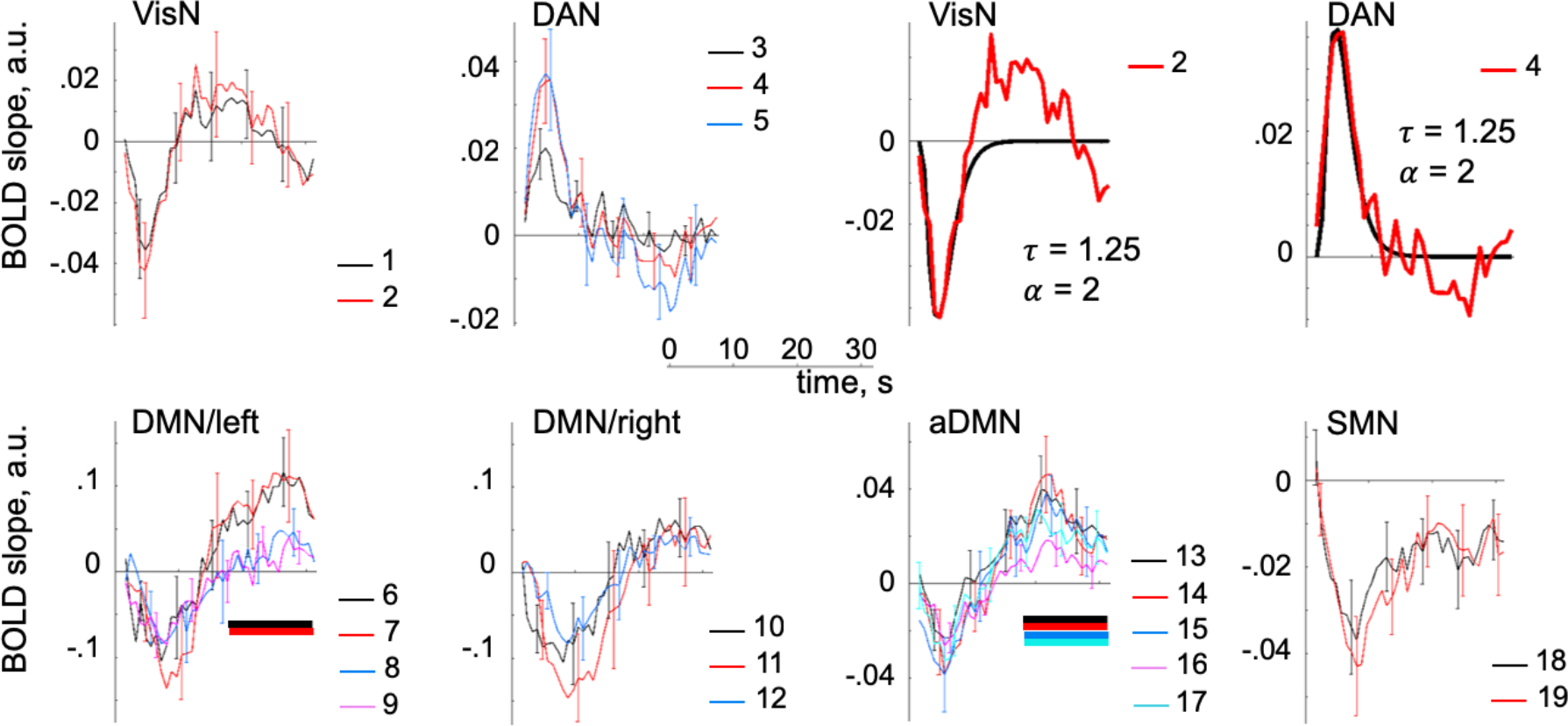
Time-courses of state-specific BOLD changes computed using a FIR approach without *a priori* assumptions about HDR shape. The slope of BOLD change per unit increase in fractional occupancy by eliciting electrophysiological network states during the trigger frames was estimated at each fMRI frame during a peri-trigger-frame time interval (time, 0 s, corresponds to the trigger fMRI frame). BOLD time-courses for HMM states were aligned relative to an 8-s pre-trigger-frame baseline and averaged across voxels in each numbered significant cluster in Figure 4 and across participants. Error bars show standard error of the group mean. The upper right panels show that VisN and DAN timecourses could be modeled by a canonical HDR function (black). Colored horizontal bars in the bottom panel indicate the time-courses that were significantly different from baseline during the 16-32-s post-trigger-frame time interval (p<0.05). HMM state name abbreviations and cluster numbers are the same as in Figures 2.

Association between lower-frequency electrophysiological amplitudes and BOLD in the DMN may be linked to an alert wakeful state^59^. In our study, the temporal pattern of electrophysiological DMN state dynamics suggested that this network might be closest to criticality -- a high energy state, when the neuronal resources are maintained at maximal availability for optimal vigilance in the uncertain world^50^. Thus, we asked whether the steep slopes of late BOLD increase per unit increase in electrophysiological fractional occupancy, which may reflect energy resources replenished to support future activity^54^, may be inherent to the DMN state of criticality.

We used the integral of the slope during the positive lobe of the DMN HDR curves (shown in Figure 5) as a metric of positive BOLD responsiveness. After the effects of age and gender were regressed out, this metric averaged across all seven DMN nodes was associated with the power law fit computed on DMN electrophysiological lifetimes (i.e., power law deviation statistic shown in Figure 3 multiplied by −1 for easier interpretation; r=0.58, p<0.05; follow-up analysis revealed that this result was driven by five nodes). Figure 6 (upper panel) illustrates associations between positive BOLD responsiveness in individual DMN nodes and the power law fit. Similar results for power law metrics computed on intervals between electrophysiological states are reported in Supplemental Figure 2. We found no association in control analyses between analogous BOLD responsiveness metrics for other HDR effects and corresponding state-lifetime-based power law statistics (abs(r)<0.33, p>0.22; see Supplemental Discussion for comments on the aDMN), except for the DAN effect (r=-0.58, p<0.05). We had no *a priori* hypothesis for the DAN state; if replicated, an association between high positive BOLD responsiveness and a DAN electrophysiological state, characterized by low amplitude (i.e., desynchronized) activity deviating from energetically optimal spontaneous patterns, would be consistent with prior evidence that the DMN and DAN represent alternate regimes of intrinsic brain function^60^.

**Figure 6.**
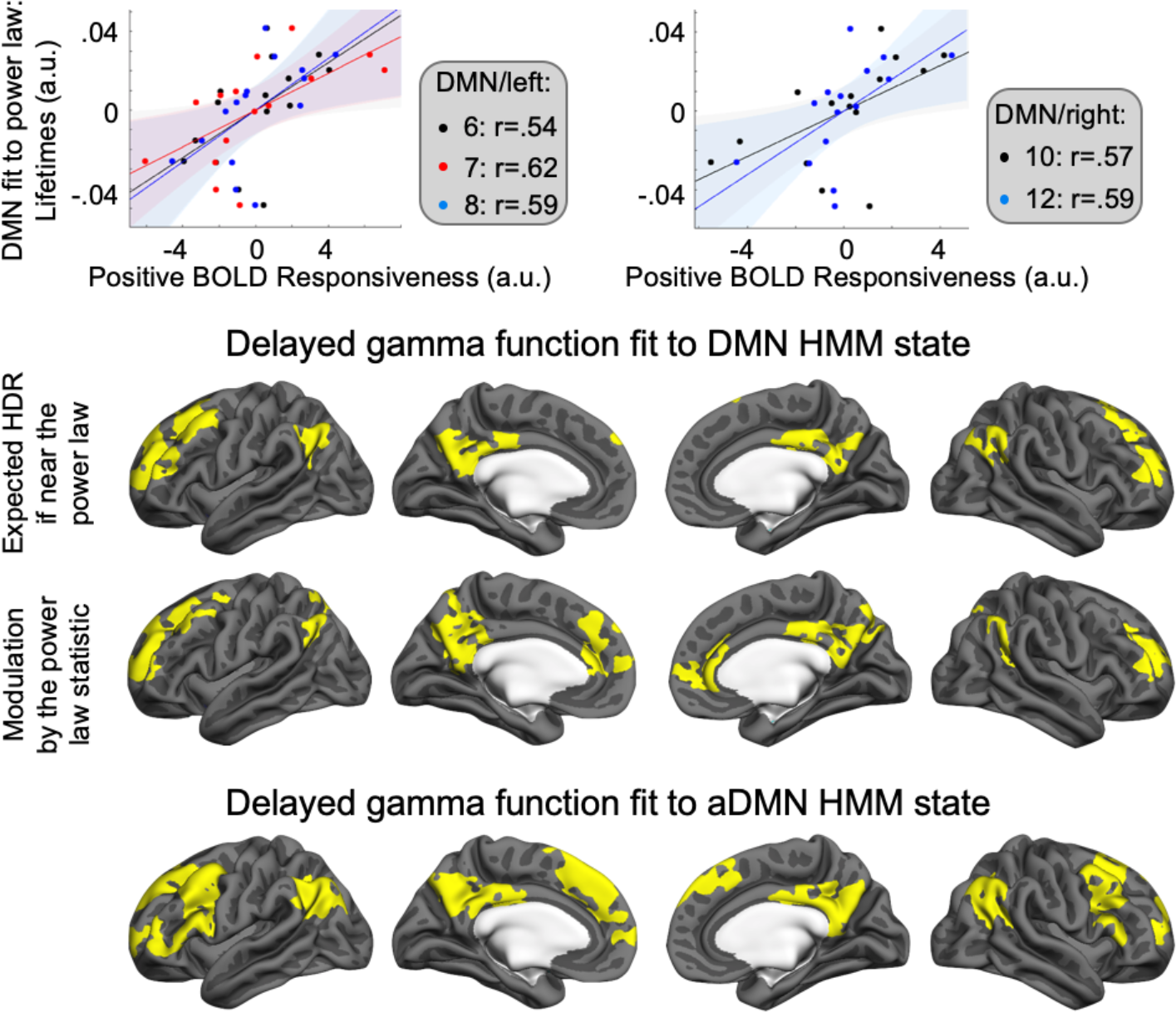
Cross-sectional association between positive BOLD responsiveness in the DMN and the temporal structure of DMN electrophysiological activity. In five numbered significant clusters from Figure 4, the integral of the slope of positive BOLD response to DMN-state fractional occupancy variation was correlated with the fit between the PDF of DMN lifetimes and the power law (negative of the deviation statistic in Figure 3); shaded areas show 95% confidence intervals; for all r-values, p<0.05 (upper panel). Significant spatial clusters (p<0.05, corrected) for the relationship between positive BOLD responsiveness and DMN lifetime-based power law fit (middle panel). Maps of positive BOLD response to DMN state if deviation of DMN lifetimes from the power law were 0 are shown in the upper row, and of an interaction between the positive BOLD effect and the DMN power law fit are shown in the lower row. Maps of positive BOLD response evoked by aDMN state (p<0.05, corrected) shown for comparison (bottom panel). The slope of BOLD change in response to frame-to-frame fractional occupancy variation was modeled with a 12-s delayed gamma response function.

We examined the spatial topography of the association between positive HDR evoked by DMN state and the power law fit for DMN electrophysiological lifetimes by modeling a voxel-wise gamma response function with a 12-s delay after state-specific electrophysiological trigger-frames (i.e., an approximate time-point when the FIR-estimated curves deviated from the fitted gamma model in our primary DMN analysis, Figure 4). In group analysis, we modelled (1) an interaction between BOLD slope (i.e., BOLD signal change per unit change in DMN state fractional occupancy) and power law fit as well as (2) expected BOLD slope if the power law deviation were 0 (i.e., the intercept). The observed activation clusters displayed in the middle panel of Figure 6 overlapped the classic DMN nodes^60^ (even if slightly different from the deactivation clusters in our primary analysis; see Supplemental Discussion). For the aDMN state, we examined the spatial topography of the late positive BOLD response. The bottom panel of Figure 6 shows activation clusters observed in a voxelwise group analysis of the BOLD slope (per unit change in aDMN state fractional occupancy) modeled with a delayed gamma response function.

### Associations with presumed cerebrovascular white-matter signal abnormalities (WMSA)

We examined possible cerebrovascular links with positive BOLD responsiveness in the DMN, previously associated with heightened energy demand^61–63^. The overall volume of WMSA evident in structural MRI data was used as a surrogate marker of general cerebrovascular decline^64–67^. After the effects of age and gender were regressed out, positive BOLD responsiveness averaged across all seven DMN nodes was associated with WMSA (r=-0.64, p<0.05; follow-up analysis revealed that this result was driven by five nodes). Figure 7A illustrates the relationship between WMSA volumes and positive BOLD responsiveness in DMN nodes. In additional analyses, correlation of WMSA with DMN electrophysiological power law fit was not significant for the lifetimes-based metric (r=-0.09, P>0.7) but was present for the intervals-based metric (Supplemental Figure 2).

**Figure 7.**
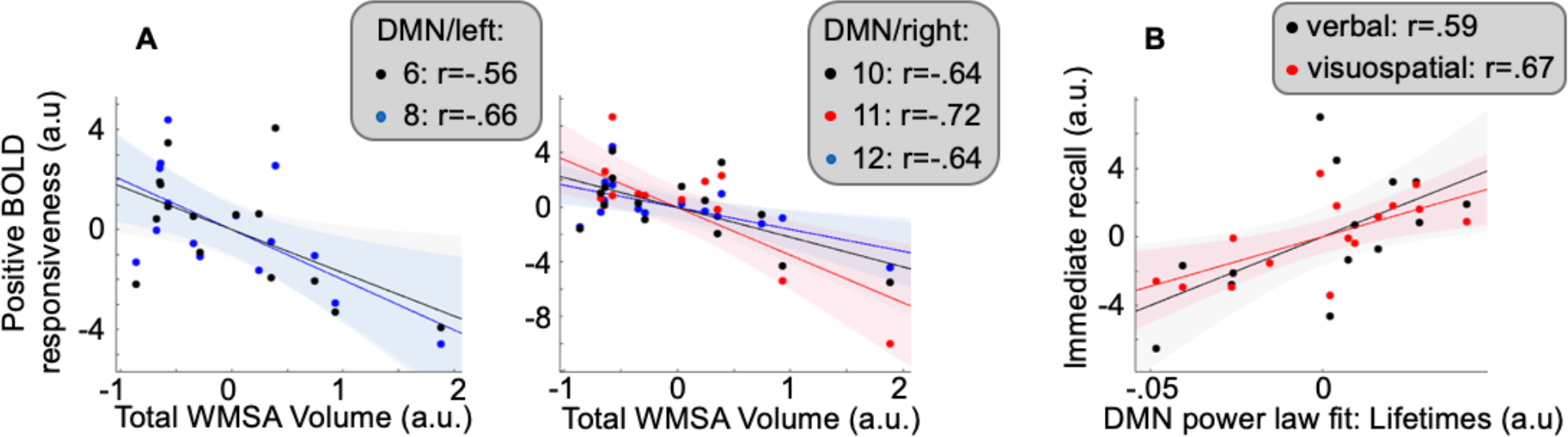
Cross-sectional associations between total WMSA volume and positive BOLD responsiveness in the DMN **(A)** and between DMN power law fit estimated based on lifetimes and immediate episodic memory recall **(B)**. Shaded areas, 95% confidence bounds; for all r-values, p<0.05. Node numbers correspond to significant clusters in Figure 4.

### Associations with memory

The DMN is consistently implicated in episodic memory^68^. Because criticality is the state of optimal cognitive readiness^69,70^, we reasoned that participants who display patterns of DMN coordinated electrophysiological activity particularly suggestive of criticality may show advantage in forming episodic memories. Indeed, scaling of neural lifetimes has been linked to behavioral parameters^46^. Figure 7B illustrates the relationship between power law fit computed based on DMN lifetimes (i.e., a metric of criticality) and performance in immediate recall trials for verbal (Wechsler Logical Memory I, WLM-I^71^) and visuospatial (Brief Visuospatial Memory Test, BVMT^72^) memories. After the effects of age and gender were regressed out, the power law fit was associated with the summed normalized WLM-I and BVMT scores (r=0.65, p<0.05; follow-up analysis confirmed the association for both verbal and visuospatial recall). These measures of memory were not associated with DMN positive BOLD responsiveness (r<0.42, p>0.12).

## Discussion

Using a novel study paradigm, EEG and fMRI data, simultaneously acquired during a resting scan, were analyzed to infer the time-courses of coordinated electrophysiological shifts in large-scale brain networks and to estimate network-specific shapes of the HDR time-locked to changing rates of electrophysiological events. An HMM approach was used to infer sequential transitions between several underlying network states in fluctuating high-density EEG source-localized to 38 regions covering the entire cortex. These transient states represented discrete spatial patterns of coordinated intrinsic electrophysiological activity lasting only a fraction of a second but recurring in a rapid, irregular succession. In a sample of older adults, inferred electrophysiological states reproduced main spatial and temporal parameters of MEG-based states previously described using similar methodology in younger^14,15^ and older adults^16^. In an event-related analysis of rapidly acquired fMRI, we leveraged the varying amounts of time spent in a given electrophysiological network state during different 800-ms recording frames. Using GLM, we deconvolved HDR patterns associated with temporal variations in fractional occupancy of electrophysiological activity in different networks. Spatially, HDR patterns observed in older adults corresponded to both maps of eliciting electrophysiological network states and classic functional networks described in prior fMRI studies^31–36^. Temporally, HDR patterns showed varying network-specific shapes ranging from predominantly monophasic to biphasic. These findings describe, for the first time, how the rapid stream of spatial coordinated changes in spontaneous electrophysiological activity relates to the morphology of local HDR—building a conceptual and methodological framework for future research on the interplay between vascular and neural dynamics in the human brain. These results also inform inferences about neural activity in older adults based on complementary information provided by EEG and fMRI.

The DMN, which is susceptible to aging and neurodegenerative disease^27^, showed electrophysiological and HDR signatures remarkably dissimilar from those of the DAN, a less vulnerable higher-order cognitive system^28,29^. The DMN functional state was characterized by increased amplitudes of 4-30-Hz band-limited electrophysiological activity, temporal parameters scaled according to a power law, and a negative-to-positive biphasic HDR. By contrast, the DAN functional state was marked by decreased electrophysiological amplitudes, temporal parameters substantially deviating from the power law, and a monophasic positive HDR. General cerebrovascular health of older non-demented adults, estimated based on signal abnormalities in structural MRI scans, predicted how steep the slopes of the positive BOLD change in response to increased coordinated electrophysiological activity were in the DMN. This positive BOLD responsiveness was associated with individual differences in the temporal structure of electrophysiological DMN state dynamics linked to memory decline. These results offer new insights into potential origins of DMN functional vulnerability in aging. In this network, which may be at the heart of neural communication across the brain^73–77^, neurovascular coupling may play a unique role in regulation of intrinsic neural activity.

The current investigation exploited high-density EEG, which may have been instrumental to attaining signal localization to cortical regions sufficiently accurate to replicate previous HMM-based pattern decomposition of MEG data^16,78^. Whereas EEG consistently achieves as high temporal resolution as MEG, the present investigation is one of a relatively small subset of studies in which high-density EEG afforded good spatial resolution^79–81^. In fact, EEG sensitivity to both radial and tangential electrical currents in the cerebral cortex–superior to the mere radial current sensitivity of MEG^82,83^–may have yielded an enhanced signal, improving HMM performance. Our analysis differentiated between two states: a DMN state with a cortical map overlapping classic DMN nodes^32,84,85^, including the precuneus, and an aDMN state that was previously linked to self-referential thought^41^. In our prior studies in younger and older adults, based on source-localized MEG, HMM did not distinguish between these two states or identify state-specific changes in the precuneus^14,16^. Only after an additional parameter—phase-locking of electrophysiological activity—was taken into consideration were similar effects detected in an MEG dataset^15^. Altogether, our data suggest that high-density EEG, even when recorded during an fMRI scan, is an effective tool for imaging rapid electrophysiological dynamics in core large-scale functional networks in older adults.

Several previous studies in younger participants show temporal associations between features of EEG and fMRI data recorded simultaneously during a resting scan^17–19,57,86^. Based on the premise that each transient electrophysiological change evokes a local HDR, these analyses show temporal correlations between BOLD time-courses as expected based on unique components of EEG activity convolved with a canonical HDR function and actual BOLD fluctuations in distinct brain networks recorded by fMRI^17–19^. Our study design built upon this evidence and took several steps forward. First, by relying on a well-validated HMM application to electrophysiological source-localized data^14–16^, we did not merely establish temporal concordance but were able to predict specific brain areas in which each electrophysiological spatiotemporal pattern would evoke an HDR. Although source-localized EEG is characterized by limited spatial resolution and inherent ambiguity due to the inverse problem^87,88^, spatial correspondence between EEG and fMRI effects was readily apparent. Second, we modeled HDR evoked by network-specific electrophysiological activity without any *a priori* assumptions about its shape. Such an analysis permitted—for the first time—quantification of the HDR timecourse for higher-order cognitive networks, which previously was not possible due to a lack of information about the neurophysiological timing of cognitive events. Third, we estimated the slope of BOLD change in response to timeframe-to-timeframe variation in fractional occupancy by electrophysiological network activity, which may be a promising metric of individual differences in neurovascular coupling. Such specialized measurement of the BOLD mechanism would be valuable for disambiguating between neurophysiological and neurovascular effects in studies of aging and neurodegenerative disorders^89^, especially since electrophysiology and fMRI have been shown to carry additive information about brain health^90^. Previously, small participant samples were sufficient to infer robust HMM states^14,16^ that were subsequently reproduced in larger datasets^91^. In the present study, we observed distinct BOLD signatures of HMM states in a relatively small participant sample. Although replication is needed, this result underscores that the HMM pattern-decomposition approach is a promising tool for developing future sensitive brain diagnostics.

In “task-positive” functional networks–generally known to show increased BOLD signal in response to cognitive loads^60^, our resting-state analysis revealed a monophasic HDR, but changes from baseline were of different polarities. Initially, we utilized a gamma response function to model BOLD responses in the entire cortex. The obtained HDR maps showed clear spatial correspondence to maps of temporally concordant electrophysiological states. To estimate the HDR time-course without *a priori* assumptions about its shape, we used FIR basis functions^23^. Previously, the shape of what is now considered a “canonical” HDR was described based on an obligatory primary sensory response in the visual cortex^20–23^. In our older adult participants, the shape of the HDR to intrinsic synchronized electrophysiological changes in the VisN and DAN was similar to the canonical shape. In the context of prior carefully designed studies showing a preserved canonical HDR in normal aging^92–94^, this finding validates our novel event-related paradigm for analysis of concurrent electrophysiological/BOLD data. In the intraparietal sulcus, which has been linked to shifts in top-down attention^95–97^, our estimate of HDR shape in older adults can benefit future fMRI studies with regard to the interpretation of the timing of evoked neural activity. Remarkably, the sign of BOLD change from baseline relative to the sign of spatiotemporally concordant change in electrophysiological amplitudes in our study is consistent with prior evidence. Previously, a negative temporal correlation was found between BOLD signal intensity and amplitude variations in lower frequency (4-30 Hz) electrophysiological activity across trials in task-related designs^98–101^ and across spontaneous fluctuations during resting-state recordings^57^. Similarly, in our study, the DAN state, characterized by reduced electrophysiological amplitudes in the 4-30 Hz band, evoked a positive HDR, whereas the VisN and SMN states, characterized by increased electrophysiological amplitudes, evoked a negative HDR.

A distinctive pattern of a biphasic HDR was evoked by DMN/aDMN electrophysiological states. Modeling the cortical HDR to each state by fitting a gamma response function was adequate to detect significant changes in regions matching classic DMN nodes^32,35^. Nonetheless, in several nodes, estimation without *a priori* shape assumptions, using a FIR basis set, revealed an initial BOLD reduction followed by a BOLD increase. In previous concurrent EEG/fMRI recordings at rest, DMN patterns were inconsistent, ranging from no electrophysiological states correlated with DMN BOLD fluctuations^18^ to separate positive and negative temporal correlations of distinct transient electrophysiological states to the BOLD signal in subsets of DMN nodes^17,19^. Such inconsistent results could occur if the modeled BOLD time-courses obtained by convolving EEG events with a monophasic HDR function were evaluated against the actual time-courses where the electrophysiological events evoked a biphasic BOLD response. The distinct HDR shapes evoked by intrinsic DMN activity may also explain limitations of the HMM in describing dynamics in fMRI-based functional connectivity^102^, where network-specific time-variable HDR properties can make state transitions appear less sharp. We consider possible reasons for the biphasic HDR in the Supplemental Discussion.

The DMN is implicated in higher-order cognition, and its nodes overlap hubs of intrinsic brain activity^74,84,103^. It is also one of the most metabolically active networks in the resting brain, showing elevated levels of cerebral blow flow (CBF)^61,62^ and glucose metabolism^63^ relative to other brain regions. As a result of this intense demand, the health of this network is likely reliant on adaptations in neurovascular interactions, such as the high energy state of criticality. At criticality, neuronal resources may be maintained at maximum availability for optimal information processing^46,50,104^, and the arising unique patterns of spontaneous activity^105^ with parameters scaled according to power law^51^ may be regulated by the pace of cellular resource replenishment^106–108^. Power law scaling was previously observed even in large-scale measurements of spontaneous electrophysiological activity from cell populations^44–47^. In our study, to describe temporal structure of activity in electrophysiologically defined networks, we estimated how much scaling of HMM state parameters deviates from power law. The DMN temporal patterns showed one of the smallest deviations from the power law among electrophysiological states. Across participants, the proximity of the DMN state lifetimes to the power law and thus, presumed criticality, predicted mental readiness to engage in episodic memory encoding as was suggested by superior immediate recall of verbal and visuospatial information. Selfregulation of the criticality regime may arise from slow built-up of energy after its dissipation by activity^109^. Besides, transient network activations may be phase-locked to ultra-slow spontaneous waves, evident in the electrophysiological and fMRI recordings, which have been suggested to reflect cortical excitability^16,110,111^. To examine the link between cerebrovascular support regulation, which may enable energy built-up, and DMN electrophysiological state patterns, we used positive BOLD responsiveness to DMN electrophysiological states. A cerebrovascular health contribution to this DMN metric was suggested by its cross-sectional inverse association with WMSA burden, a surrogate marker of general cerebrovascular decline^112^. Our results revealed a strong association of positive BOLD responsiveness in DMN nodes with the DMN power law fit metrics for inter-state intervals and state lifetimes that—in whole-cortex-wise analysis— was confirmed to selectively localize to the DMN network. Thus, HDR regulation to variable rates of spontaneous electrophysiological activity in the DMN may be linked to sustaining the optimal functional regime near criticality.

Overall, our findings suggest that classic functional brain networks are characterized not only by their spontaneous neural activity patterns, but also by their HDR shape signatures. Remarkably, HDR shapes were highly similar within functional networks, even when considerable differences existed between networks. In future research, it will be important to gain a better understanding of how the HDR, which faithfully follows coordinated changes in spontaneous electrophysiological activity, might adapt to the properties of unique neural networks. Interestingly, it has been shown that slower timescales of neural activity may characterize brain regions with larger numbers of connections^113^, perhaps because such regions may integrate more information. It may be that HDR has adapted to optimally support such functional differences among networks. Our study describes the BOLD response to changes in the rate of network-specific lower-frequency electrophysiological activity but was not designed to measure high-frequency gamma bursts (30-90 Hz). Methodological difficulties in discerning gamma activity in the presence of high frequency recording noise can be overcome by employing interleaved EEG/fMRI designs^114^ and specialized HMM analyses^115^. Gamma activity is linked to high metabolic demand^116^ and may impact how unique patterns of cross-frequency^117–121^ and cross-cortical-layer^114,122–124^ integration during electrophysiological network states are coupled with the HDR. Another methodological advance would involve concurrent EEG recording within BOLD/CBF measurement designs that can estimate oxygen consumption^125^. In our study, electrophysiological states in sensory/motor networks evoked negative-going BOLD responses. Although the magnitude of these responses did not relate to overall temporal electrophysiological patterns as quantified by the power law fit of their parameter PDFs, it is interesting to consider a potential role of such activity states in reducing the cost of brain function during resting state. For instance, with limited sensory input during rest, sensory networks may undergo brief states of lowered metabolic demand (i.e., simultaneous reductions in oxygen consumption and CBF)^126^ or reduced metabolic-vascular coupling (i.e., reduced CBF relative to oxygen consumption)^127^. Our study used short (800-ms) BOLD recording frames to capture the fastest neurovascular dynamics possible while safely recording EEG/fMRI in older participants. Nonetheless, electrophysiological HMM states exhibit a slower temporal structure (i.e., quasiperiodic reoccurrence rate fluctuations <0.1 Hz)^14,16^, where transient states of coordinated network activity are embedded into the phase of ultra-slow fluctuations, likely, related to changing cortical excitability^111^. Simultaneous BOLD/CBF designs can take advantage of this slow structure while recording data in longer (2-s) frames.

In summary, in a novel analysis of concurrent EEG/fMRI, we observed a readily apparent spatiotemporal correspondence between electrophysiological and hemodynamic fluctuations in spontaneous brain activity in older adults. The brain at rest exhibited an ongoing stream of network-specific electrophysiological changes, with each network displaying unique features of neurovascular coupling. Whereas VisN and DAN electrophysiological states evoked a near-canonical HDR shape, establishing the validity of our analysis, the DMN state showed a unique negative-to-positive biphasic HDR. Importantly, lower positive BOLD responsiveness in the DMN across our sample of older adults was predicted by deteriorating cerebrovascular health and associated with disruptions in the natural temporal pattern of DMN electrophysiological activity linked to memory decline. If future work verifies that temporal parameters of fast natural activity, which may be protective against Alzheimer’s pathology^128^, are regulated by neurometabolic or neurovascular coupling, it may be possible to deliver preventive treatments through innovative technologies such as near-infrared light^129^. Future research delineating diagnostic procedures for HDR dysregulation could lead to the design of optimal light treatment schedules to enhance mitochondrial and vascular function by targeting specific layers and cell types in the cerebral cortex^114,130^.

## Methods

### Participants

Fifteen older adults were recruited to take part in this study through the Brain Aging and Dementia Laboratory at Massachusetts General Hospital from a local longitudinal cohort or through community outreach. Individuals were non-demented with Mini-Mental State Examination scores >24. Because we primarily aimed to study DMN and DAN, we estimated the sample size we would need based on previous findings of a rather large effect size (Cohen’s d between 0.7-0.9; N.B., d=t/√n) in fMRI-based within-subject comparisons in these networks^131,132^. Because we aimed to study the fMRI signal associated with changes in the electrophysiological neural activity, we expected smaller error variance than in prior studies where the measurement error might have been influenced by trial-to-trial variation in the neural response to cognitive stimulation. Thus, we aimed to detect an effect size of at least 0.8 (one-sample, two-tailed t-test), which would require 15 participants to have 80% power or greater at a 0.05 level of significance^133^. Additionally, we aimed to test targeted hypotheses about specific cross-sectional associations (e.g., between the neurally-evoked HDR and a surrogate marker of the global cerebrovascular health). We anticipated that we would have 80% power to detect a strong correlation of 0.61 or greater at a 0.05 level of significance (one-tailed) with 15 participants^134^. We excluded participants for significant health concerns that would prevent participation or would be likely to confound study results. These conditions included major neurological or psychiatric disorders (e.g., Parkinson’s disease, Huntington’s disease, vascular dementia, clinical stroke, brain surgery, psychosis, severe major depression, and moderate to severe traumatic brain injury) or any substantial systemic illness. All individuals had at least a high school education (i.e., 12 years).

The local Institutional Review Board approved this project, and informed consent was obtained from each participant. Participant demographic characteristics and cognitive/functional assessment scores are provided in Table 1.

**Table 1.** Demographic and clinical characteristics of study participants.

| Assessment | Mean±std |
| --- | --- |
| Age | 72.60±6.96 |
| Gender | 7 males |
| Years of education | 15.80±3.00 |
| MMSE | 27.87±1.68 |
| MoCA | 24.13±3.91 |

Note: MMSE, Mini-Mental State Examination; MoCA, Montreal Cognitive Assessment, std, standard deviation.

### Data acquisition

Imaging was conducted with a Siemens Prisma 3-Tesla whole-body MR scanner (Siemens Medical Systems) using a 32-channel head coil. High-resolution scanning protocols developed by the Human Connectome Project^135^ were employed in both structural MRI (T1-weighted multi-echo magnetization-prepared rapid gradient echo; voxel size, 0.8 mm^3^; field of view, 256×240×166; TR/TI=2500/1000; TE=1.8/3.6/5.4/7.2; flip angle, 8°) and fMRI (two 10-min-long consecutive resting-state T2*-weighted scans sensitive to BOLD contrast: 2.0-mm isotropic voxels covering the whole brain [72 oblique-axial slices] were acquired with a 2D multiband [MB] gradient-recalled echo echo-planar imaging sequence; MB8; TR/TE=800/37 ms; flip angle, 52°). EEG (filter, 0.01-200 Hz; sampling rate, 1000 Hz) from a 256-channel net included in an MR-compatible Geodesic 410 MR System (Electrical Geodesics, Inc., [EGI], a Philips company) was acquired concurrently with fMRI. EEG sampling was temporally aligned to MR gradient artifacts using the scanner’s 10-MHz clock, and triggers marking each fMRI recording frame, and thus artifact onsets, were recorded by the EEG system. EEG channels directly below and to the side of each eye were used to track the electrooculogram to identify epochs containing artifacts due to vertical and horizontal eye movements or blinks. For ballistocardiogram artifact removal, electrocardiogram (ECG) was acquired using two MR-compatible ECG leads (InVivo Corp.) placed on the chest. Participants were instructed to keep their eyes open during resting functional scans and refrain from motion. Structural images of each participant were registered to the MNI152 standard brain to allow subsequent EEG source space analyses in MNI space. Locations of EEG sensors with respect to anatomy were determined by registering head surface sensor positions, digitized using GPS Photogrammetry (EGI), to the head surface extracted from structural MRI.

### Analysis tools

Software packages used in data analysis included FSL^136^, SPM12^137^, Fieldtrip^138^, AFNI (https://afni.nimh.nih.gov/), and FreeSurfer^139,140^. Sample MATLAB (The MathWorks Inc., Natick, MA) scripts utilized for EEG analysis can be found at https://www.psych.ox.ac.uk/research/ohba-analysis-group. Scripts used for event-related fMRI analysis are included in the FreeSurfer Functional Analysis Stream - FSFAST (https://surfer.nmr.mgh.harvard.edu/fswiki/FsFast).

### Pre-processing of EEG data

Gradient switching artifacts were removed using a 10-sample moving average subtraction, and ballistocardiogram artifacts were reduced using optimal basis sets^141^ implemented in NetStation (Electrical Geodesics, Inc.). Each EEG recording was then visually inspected to identify channels and/or time-intervals of data containing obvious artifacts (e.g., extremely high variance), which were discarded. Residual cardiac and ocular artifacts were removed by Independent Component Analysis (ICA). Following artifact cleaning, EEG data were converted to SPM12^137^, frequency filtered into a 4-30 Hz band— carrying most information about brain heath^142^ and characterized by a relatively high signal-to-noise ratio in electrophysiological recordings^143^, and downsampled to 200 Hz.

### Electrophysiological time-courses in the source space

Forward modeling used a Boundary Element Model^144^ with scalp, skull, and brain surface reconstructions. A linearly constrained minimum variance scalar beamformer, implemented in SPM12, was used to project band-limited EEG data onto a regular 8-mm grid spanning the entire brain^145–147^. Beamforming provides verifiable neural source estimates and effectively rejects interference from non-brain sources in the electrophysiological signal^148,149^. Projected data were scaled by an estimate of projected noise to account for variations in the sensitivity of the beamformer at different brain locations^145,146^.

Thirty-eight anatomical regions of interest (ROIs) were selected based on a group spatial ICA of fMRI resting recordings from the first 200 participants in the Human Connectome Project database^150^. This same set of ROIs was previously used in network analyses based on resting state MEG^30,151^. Regional time-courses were obtained based on projected band-limited EEG data using principal component analysis of voxel time-courses within each ROI, normalized so that the positive peak had a height of unity in all regions. The timecourse for an ROI was represented by coefficients of the principal component accounting for the most variance, weighted by the strength of the ICA spatial map. Activity with zero phase-lag, which likely contains “signal/spatial leakage” to multiple EEG sensors and may lead to inflated connectivity estimates, was accounted for by symmetrically orthogonalizing all ROI time-courses simultaneously^30^. Multivariate symmetric orthogonalization^152,153^ produces a unique solution that is unaffected by reordering of ROIs and constitutes an optimal set of mutually orthogonal ROI time-courses that are minimally displaced from the uncorrected original time-courses (as measured by least-squares distance). By being multivariate, this method can also account for any spurious associations inherited from true connections, also known as “ghost interactions”^30,154^.

Following leakage reduction, the amplitude envelope of the electrophysiological band-limited activity for each ROI time-course was derived by taking the absolute value of its Hilbert transform and downsampling to 40 Hz^155^. These time-courses of oscillatory amplitude fluctuations were demeaned and normalized by global (i.e., over all voxels) variance and then concatenated temporally across all participants.

### HMM

Utility of HMM in describing the neural network dynamics has been highlighted^16^, and HMM inference was conducted using previously described computations^14^. An HMM with 10 states was inferred based on the time-courses of band-limited electrophysiological amplitudes from 38 ROIs concatenated across all participants. Previously, we successfully used a 10-state HMM in a sample of older adults^16^, and in an HMM analysis comparing models with 3-12 states, free energy estimates exhibited a floor effect at 10 states^156^. Additional HMMs with 12 and 14 states yielded similar core results.

The HMM infers transient states in which coordinated changes in electrophysiological amplitude recur in a distinct set of brain regions as well as a time-course for each state. Individual HMM states were defined by a unique multivariate normal distribution over the ROIs (i.e., mean vector [M×1] and covariance matrix [M×M], where M=38, the number of ROIs)^157^. Previously, we showed that this relatively simple observation model of multiple HMM states can describe rich spatiotemporal dynamics in the living brain^16^. To account for variability in HMM inference due to different initializations, 10 realizations were performed, and the model with the lowest free energy was selected. The Viterbi algorithm was applied to identify the most probable state at a given time point^157^. Time-courses for each state were constructed as indicator variables specifying time intervals when the state is most probable.

### Spatial maps of electrophysiological states

Anatomical regions exhibiting state-specific activity (i.e., changes in electrophysiological amplitudes during an HMM state relative to the average over time) were mapped by computing partial correlations within the GLM framework^14^. In all GLM analyses, we employed a design matrix (T×K), where K is the number of states and each of the K columns is a state time-course with T time-points^158,159^. For each participant at each of the 38 *a priori* ROIs, multiple linear regression was performed with the time-course of electrophysiological activity as the dependent variable. Prior to GLM, both the design matrix and ROI data were normalized to have zero mean and unit variance. Estimates of partial correlation coefficients between each state and ROI data yielded a set of K spatial maps. These maps were averaged across participants and visualized on the cerebral cortex.

### Temporal dynamics of electrophysiologically defined networks

For the purpose of computing temporal metrics, states were classified as being on or off by choosing the most probable state at each time point (i.e., Viterbi path). Lifetimes of inferred HMM states were defined as the amounts of time spent in each state before transitioning out of that state. Intervals between network activity states were defined as the amounts of time between consecutive occurrences of a given HMM state. A PDF was used to describe likelihood of different lifetime/interval values during the recording period. To quantify temporal structure of coordinated activity in each network, we computed a Kolmogorov-Smirnov (KS) statistic indicating the degree to which a PDF of the lifetimes in each state deviates from a modeled power law distribution. Similar metrics were also obtained for intervals between network activity states. Specifically, the KS statistic reflects the difference between the empirical PDF and a model power law PDF fitted to the data using maximum likelihood estimation^52^. To examine differences between DMN/aDMN states and the comparison DAN state, we used an omnibus rmANOVA with state as a factor (DMN, aDMN, and DAN) and deviation from power law as a dependent variable, while regressing out the effects of age and gender. Huynh and Feldt correction was applied to repeated measures with >1 degree of freedom^160^, and the effect size in follow-up simple comparisons was quantified using Cohen’s d^161^.

### Pre-processing of fMRI data

The first ten volumes of each functional run were discarded to allow for T1-equilibration effects. To correct for motion, all volumes in each run were aligned to a single volume chosen from the middle of the run using the AFNI algorithm. For each participant, these middle-run volumes were then aligned to 3D structural images using a combination of automated procedures from FSL/FLIRT and FreeSurfer/bbregister. Brain mask was created using FSL’s BET. Structural images were also used to construct two-dimensional (2D) models of individual cortical surfaces using FreeSurfer algorithms. Both 3D and 2D structural models were registered to an average template—a model of brain volume and surface based on an independent sample of 40 adults registered to the Montreal Neurological Institute atlas (MNI305). Intensity-normalized functional time series were resampled to the surface-based average template and smoothed with a 5-mm full width half-maximum Gaussian kernel.

### Maps and time-courses of HDR evoked by electrophysiological network states

FMRI data were analyzed under GLM assumptions. Our aim was (1) to identify voxels where the BOLD signal was modulated following changes in the occurrence rate of each electrophysiological HMM state and (2) to describe the state-specific shapes of such BOLD signal changes. In principle, the fMRI analysis was similar to that used to derive electrophysiological spatial maps of HMM states. However, state time-courses were modified by computing fractional occupancy (i.e. proportion of time spent) in a given electrophysiological state during each 800-ms fMRI recording frame (Figure 1). We used two different approaches to estimate the state-specific HDR. First, the BOLD response, following trigger timeframes with non-zero fractional occupancy by each HMM state, was modeled at each cortical voxel using a gamma response function. Participant-specific statistical models removed low-frequency drifts in the data, regressed out head motion artifacts, and estimated the slope of BOLD change in response to changes in fractional occupancy by each state (parametric modulation by a continuous variable). Group statistics were computed treating participants as a random effect, regressing out the effects of age and gender, and correcting for multiple comparisons using simulation testing. We determined the likelihood that an activation cluster of the observed size (p<0.05, uncorrected, in each adjacent voxel) could occur by chance by running 10,000 Monte Carlo simulations of synthesized white Gaussian noise using parameters of the functional analysis. We report Cohen’s d ranges across voxels in each network, while excluding outliers in the top/bottom 10% of voxels. The second approach included an ROI analysis employing a FIR model. For each participant, unbiased estimates of BOLD signal slope at 50 consecutive MRI frames time-locked to trigger frames with non-zero fractional occupancy by each electrophysiological state were obtained without making *a priori* assumptions about HDR shape. For each ROI, we calculated a BOLD slope time-course by averaging across included voxels at each time-point.

These two analysis approaches were applied iteratively. Our initial whole-cortex analysis was based on a canonical gamma response function (*α*=2, *τ*=1.25). For each significant cluster observed in this whole-cortex analysis, we defined a circular ROI (5-mm diameter) around the peak BOLD change and estimated the state-specific BOLD response time-courses in these ROIs by applying the FIR approach. The optimal gamma response function, fitting each BOLD time-course, was determined using maximum likelihood estimation, and the revised gamma function parameters (*α, τ*) were used to generate final cortical maps of the state-specific HDR. Significant clusters (p<0.05, corrected, while taking into account initial and final gamma fit analyses) were displayed on the inflated average template of the cortical surface. These significant clusters in their entirety were used as ROIs to estimate final state-specific BOLD time-courses in the FIR approach. To estimate how similar/different the optimal gamma function parameters were between the nodes within/between different networks, we used an omnibus rmANOVA with a factor of within vs. between networks and the mean Euclidean pairwise distances between (*α, τ*) points in Euclidean space, computed for each participant, as a dependent variable, while regressing out the effects of age and gender. Additionally, for VisN and DAN nodes, where a canonical gamma response appears a suitable model, we computed a Kolmogorov-Smirnov (KS) statistic indicating the degree to which the BOLD time-courses of each participant deviate from the fitted shape-flexible and canonical gamma models. We examined if the amount of deviation was different between the models, in rmANOVAs for each node with a factor of optimal vs. canonical gamma function, while regressing out the effects of age and gender.

### State-specific late HDR

To quantify the late HDR change associated with each HMM state, we computed the integral of the FIR-estimated BOLD slope within a time-window of 16-32 s after the electrophysiological trigger frames. Given prior evidence of a positive correlation between low-frequency electrophysiological activity and BOLD signal intensity in the DMN^57,58,162^, a BOLD signal increase was expected to be evoked by DMN and aDMN HMM states. We tested the statistical significance of the late positive HDR evoked by these states in an omnibus rmANOVA that examined the main effect of change from baseline during the late epoch and its interaction with the ROI factor (defined based on DMN and aDMN state-specific significant clusters in whole-cortex analysis), while regressing out the effects of age and gender. Huynh and Feldt correction was applied to all repeated measures with >1 degree of freedom^160^. In follow-up analyses, we examined simple effects of the late BOLD change from baseline in each ROI. In a similar control rmANOVA, we checked for any significant BOLD effects in the late time-window for DAN, VisN, and SMN states (ROI factor defined based on the clusters specific to these states).

To understand the functional significance of the late positive HDR in the DMN, we considered whether this upregulation in HDR was related to sustenance of the overall temporal pattern of DMN coordinated electrophysiological activity. In cross-sectional analysis, we again used the integral of the FIR-estimated BOLD slope to quantify the DMN HDR positive lobe (a metric of positive BOLD responsiveness) and computed a partial cross-sectional correlation, regressing out the effects of age and gender, between the mean of these metrics across DMN state-specific ROIs and the power law fit of the electrophysiological PDFs of DMN state lifetimes and intervals (for easier interpretation, we used “fit” computed as the KS deviation statistic multiplied by −1). In follow-up analyses, we computed additional partial correlations to determine which DMN ROIs contributed to the association between positive BOLD responsiveness and the electrophysiological power law fit and visualized the associations using scatterplots. In similar control analyses, we checked for associations between BOLD responsiveness metrics in other networks (i.e., the integral of the FIR-estimated BOLD slope in the early HDR lobe for all states and in the late lobe for the aDMN) and the corresponding power law fit of state-specific electrophysiological lifetimes/intervals.

To describe the cortical topography of late positive HDR effects, we used a whole-cortex analysis approach. We modeled the HDR time-locked to the electrophysiological trigger frames but fitted a gamma response function with a delay to capture the late epoch. Participant-specific statistical models were computed as described above. For the DMN state, group analysis modeled an interaction between the BOLD parameter and the power law fit of electrophysiological lifetimes as well as the expected BOLD parameter if the deviation from the power law were equal to 0 (i.e., the intercept). For the aDMN state, group statistics were computed for the main effect of the BOLD metric change from baseline. Significant clusters (p<0.05, corrected) were displayed on the inflated average template of the cortical surface.

### Associations with WMSA

Structural MRI images were processed using the standard recon-all stream in FreeSurfer, and reconstructed brain volumes underwent automatic segmentation of grey and white matter. Volumetric measurement of WMSA, a surrogate marker of cerebrovascular damage^64–66^, was performed using a FreeSurfer-based validated tool^67^, and WMSA volumes were adjusted to control for total intracranial volume differences and natural log transformed due to their non-normal distribution. In cross-sectional analysis, we examined a possible link with cerebrovascular health by computing a partial correlation, regressing out the effects of age and gender, between WMSA volumes and positive BOLD responsiveness averaged across DMN state-specific ROIs. In follow-up analyses, we computed additional partial correlations to identify the DMN ROIs contributing to the association between WMSA volumes and BOLD responsiveness.

### Associations with memory

Readiness to engage in episodic memory encoding would be expected to influence capacity for immediate recall, which is associated with neural activity patterns in the DMN dissociable from those linked to delayed recall^68^. Immediate verbal recall was assessed using the WLM-I^71^, which orally presents two short stories (one story twice) and asks participants to retell each story from memory immediately after hearing it. Immediate non-verbal recall was evaluated with the BVMT^72^, which consists of three trials, each presenting six geometric designs on a sheet of paper, and asks participants to reproduce the designs from memory immediately after each trial. Previously, scaling of neural lifetimes has been linked to behavioral parameters^46^. We examined whether the power law fit of the DMN PDF of electrophysiological lifetimes, suggestive of a mental state of optimal readiness to engage in information processing (i.e., criticality)^69,70^, predicts immediate episodic recall. In cross-sectional analysis, we computed partial correlations, regressing out the effects of age and gender, between the power law fit metric and the sum of normalized scores on the WLM-I and BVMT. In follow-up analyses, we used additional partial correlations to examine contributions from verbal and non-verbal recall to this association.

## Code availability

Code is available from the authors upon request.

## Data availability

Data is available from the authors in accordance with procedures approved by the Mass General Brigham Institutional Review Board.

## Author Contributions

T.S. and D.H.S. designed the study. T.S., M.W.W. and J.W.H. developed the method. T.S. and J.W.H. performed analysis. T.S., J.W.H., C.M.H., and K.A.S. conducted experiments. T.S. and J.W.H wrote and revised the manuscript. M.W.W. and D.H.S. reviewed and edited the manuscript.

## Acknowledgements

This research was carried out in whole or in part at the Athinoula A. Martinos Center for Biomedical Imaging at the Massachusetts General Hospital.

## Supplemental Information

### Supplemental Discussion

#### Biphasic DMN HDR: Spectral properties of neural activity

The DMN has long been known to show unique BOLD features, including well-known reductions evoked by sensory loads^163^. One specific distinction of the DMN that could account for the biphasic negative-to-positive HDR may be in how the BOLD signal is influenced by spectral properties of neural activity. Distinct neuron-type-specific mechanisms that give rise to synchronized activity in the gamma band (~30-90 Hz) have been demonstrated to contribute to BOLD changes independently from those in lower-frequency bands^99,164^. Even though the gamma-BOLD relationship may be similar across the DMN and other networks, the BOLD response to the activity in the lower frequencies in the DMN may be distinctive.

The gamma activity patterns, coupled to neuronal spiking^165–169^ and dependent on fast spiking interneurons^164^, are thought to represent information content of cognitive processes^170,171^. Increases in such fast neural activity are associated with BOLD rises, which are consistent across various neural networks^110,162,172–179^. In the current study, gamma was not quantified due to relatively low signal-to-noise ratio in this frequency band in our EEG recordings. However, an observation that amplitudes of lower-frequency electrophysiological activity (4-30 Hz), previously implicated in controlling neuronal spiking^180^ and the gamma through inhibition^117–120^, were increased during the DMN state, as well as the VisN/SMN states, suggested that the gamma activity might have undergone transient suppressions. Reductions in the fast neural activity might contribute to the negative lobe of the BOLD response evoked by these states. Transient modulation of fast activity during resting state may serve to refresh the information content of network patterns^169,180,181^, promoting memory replays of previously learned information^181^ as well as exhaustive “exploration” of neural activity patterns to maximize responsiveness to environment^70^.

The neuronal mechanisms underlying the lower-frequency activity (~4-30 Hz) may be different between the DMN and VisN/SMN states and may be associated with differing energy demand. In the visual and sensorimotor cortices, involvement of a specific type of inhibitory interneurons that control vasoconstriction^25^ may underlie the temporal correlation between increased low-frequency electrophysiological amplitudes and decreased BOLD intensity^57,98–101,162^. In contrast, in the DMN, a positive correlation has been observed in monkeys between electrophysiological activity at lower frequencies and polarographic oxygen measurements, which are analogous to the BOLD signal^162^. Similarly, in the human brain, lower-frequency electrophysiological activity in the DMN has been linked to the positive BOLD response^57,58^. Because the activity amplitudes are especially high in the DMN electrophysiological state^15^, high energy demands of mounting a synchronized inhibitory drive in this network may account for absence of a vasoconstricting response. In the present study, the positive BOLD response to the DMN state was observed in the later epoch, likely occurring after the eliciting neural activity was over and replenishing cellular resources needed for future activity events^54^. This may explain why responsiveness of the positive BOLD predicted the overall temporal pattern of the DMN states recurrence.

Notably, the timing of the late BOLD response to low-frequency electrophysiological changes in our study was consistent with prior clues in the literature. Previously, BOLD responses to task-evoked changes in low-frequency electrophysiological activity in the DMN were prolonged or observed with delays (~10-s long)^184^. In our study, the biphasic negative-to-positive BOLD in the DMN was likely due to summation between an earlier negative response to transient suppressions of high-frequency neural activity and a prolonged/later positive response to the increase in lower-frequency electrophysiological activity. In fact, a biphasic BOLD response in the DMN—but in the reverse positive-to-negative direction—was observed during a relational learning task^185^. It is possible that opposite modes of cross-frequency dynamics in the DMN during rest and a learning task^186^ produce a biphasic response but with reverse directions. In a similar vein, longer HDR delays to task-related lower-frequency electrophysiological activity in the SMN^187^, but not VisN^164^ are consistent with the trend evident in our study toward prolonged BOLD effect to the SMN state (Figure 5; even though not significant, the SMN time-course remained lower than baseline in the late epoch) but not VisN state.

#### Biphasic DMN HDR: Metabolic-vascular coupling

Alternatively, unique features of metabolic-vascular coupling might underlie the biphasic DMN HDR. Previously, when the DMN was compared to frontoparietal control and primary sensorimotor networks, its negative BOLD response to stimulation was characterized by an increased ratio between the cerebral metabolic rate of oxygen consumption and CBF^125^. This suggests a reduced responsiveness of CBF, perhaps due to distinct vascular biomechanics or regulation by the sympathetic nervous system^188^. If, in our study, such reduced coupling contributed to the initial negative BOLD evoked by the DMN electrophysiological state, then a delayed positive BOLD response might be expected when activity-induced pressure is relieved and coupling is normalized^127^. Another relevant piece of prior evidence is that aerobic glycolysis during rest is increased in the DMN relative to other brain regions^189^. It is possible that DMN neural activity triggers astrocytes to switch to aerobic glycolysis (i.e., energy production without oxygen), which would permit neurons to utilize all available tissue oxygen^26^. Therefore, only after an initial period of optimized oxygen consumption, marked by increasing tissue deoxygenation and a negative BOLD response, would an eventually needed blood influx generate a positive BOLD effect. Interestingly, vascular blood can also be “stolen” across adjacent cortical regions^190^. In the DMN, functional subspecialization is detected between adjacent subdivisions within each node^77^, which might explain why we observed slight loci differences for negative and positive DMN BOLD changes. It may be that redistribution of blood is optimized to support functional requirements of anatomical subdivisions.

#### Neurovascular dynamics of the aDMN

DMN and aDMN states showed similar patterns of biphasic HDR, which was not surprising given their spatial overlap in coordinated activity. Nonetheless, in contrast with the neurovascular dynamics of the DMN state, there was no significant association between positive BOLD responsiveness and temporal patterns of electrophysiological activity (quantified by a power law fit) in the aDMN state. One possible explanation for this distinction could be that the aDMN state corresponds to self-referential thought. Establishing and buffering internal trains of thought from disruptions by the external world may involve intricate cognitive-control neural dynamics, including inhibitory influences both among the inferior frontal gyrus and DMN medial nodes^41^ as well as from the frontoparietal control network^191,192^. Consistent with such complex dynamics, the aDMN state evoked initial BOLD reductions involving the medial and inferolateral prefrontal cortices, followed by a positive BOLD response encompassing the dorsolateral prefrontal and medial parietal cortices. Our HMM approach used a simple observation model based on lower-frequency electrophysiological activity amplitudes. Thus, even if able to distinguish a DMN state from other states, this analysis alone would not comprehensively describe the underlying neurophysiological processes. On the contrary, the evoked HDR (estimated using a combination of the HMM and information from the EEG and BOLD data) would be expected to more fully represent underlying neural changes, including inhibitory neuronal activity, which can result in a positive BOLD response^193,194^. Engagement of cognitive control by the aDMN state would explain why temporal activation patterns may be less regulated by the pace of replenished metabolic resources (i.e., BOLD responsiveness) and why lifetimes of the aDMN state show increased deviations from scaling according to a power law.

## Supplemental Figures

**Supplemental Figure 1.**
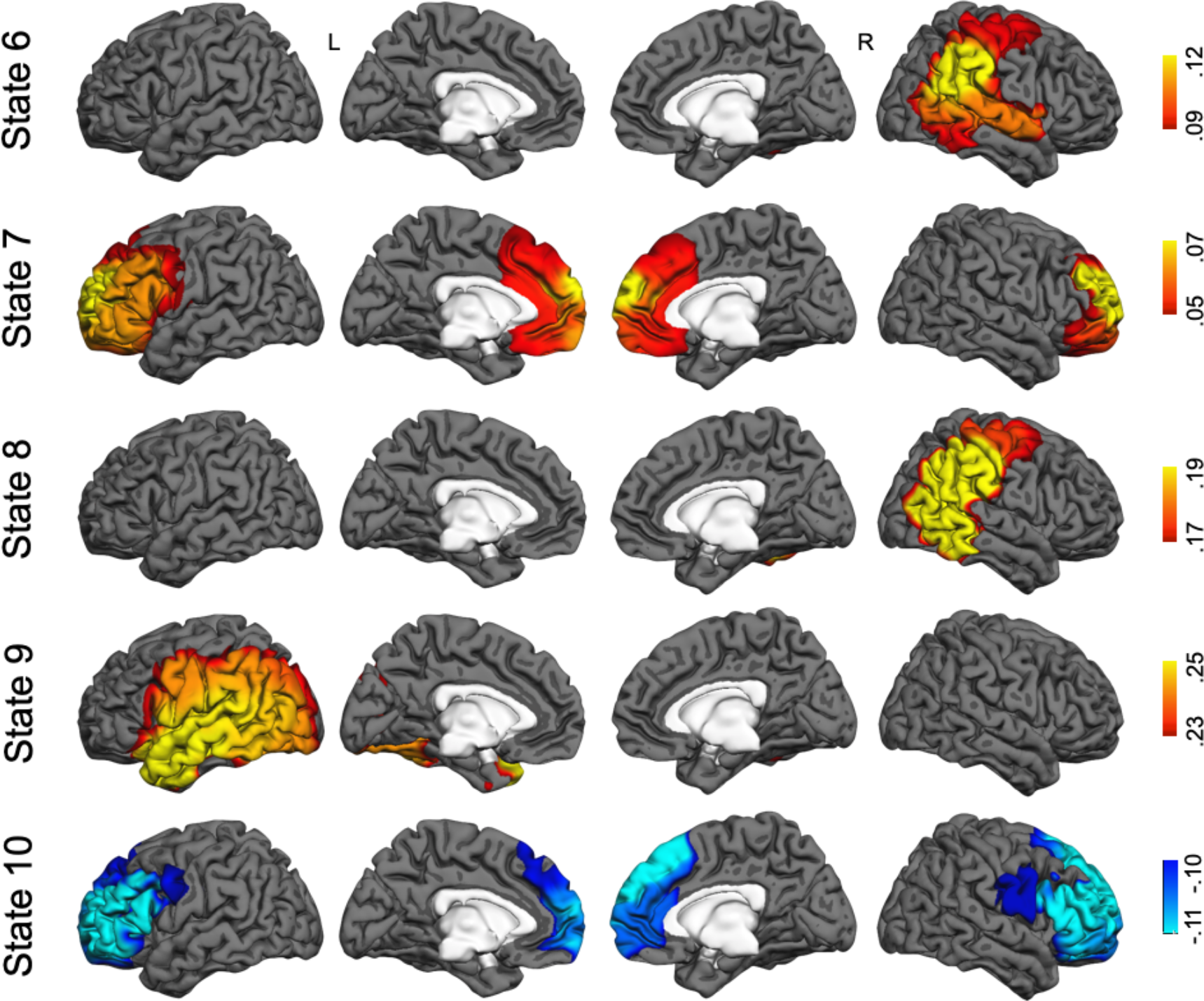
Maps of state-specific increases (in yellow/red/brown) and decreases (in blue) in electrophysiological amplitudes during five additional network states inferred by the HMM with 10 states. Each map shows partial correlations between the state time-course and the electrophysiological amplitudes in 38 ROIs. Investigating the HDR to these states was beyond the scope of this study and will be the matter of future research.

**Supplemental Figure 2.**
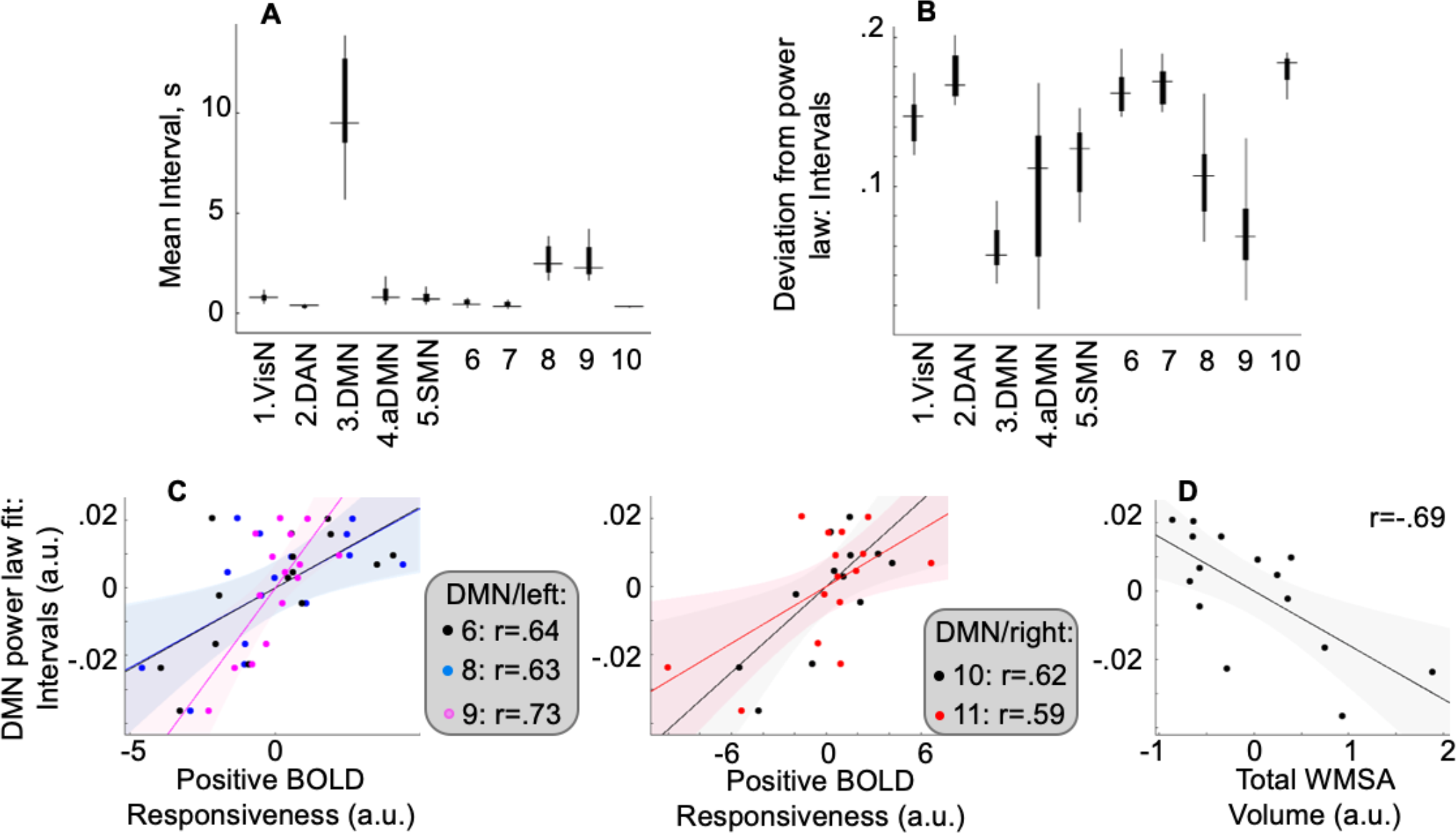
Upper panel: The temporal structure of intervals between the inferred HMM states. For each electrophysiologically defined network and participant, we computed the mean of intervals between coordinated activity states **(A)** and the deviation between empirical interval data and the power law function fitted using maximum likelihood estimation **(B)**. Boxplots for the five salient states (1–5; shown in Figure 2) and five other modeled states (6–10; shown in Supplemental Figure 1) summarize the group results (medians and 25th/75th percentiles; whiskers extend to the most extreme points not including outliers). A comparison of DMN/aDMN states to the DAN state based on the PDF of intervals between network activity states revealed differences in the extent of deviation from the power law (effect of state in omnibus ANOVA, F=49.73, p<0.05; follow-up: DMN<DAN, F=735.74, Cohen’s d=5.15, DMN<aDMN, F=8.43, d=.68, aDMN<DAN, F=28.12, d=1.45, ps<0.05). Thus, the interval parameter, similar to the lifetime parameter, suggested that the DMN state shows temporal activity structure closest to critical behavior. Lower panel: Cross-sectional association between cerebrovascular factors and the temporal structure of intervals between DMN electrophysiological activity states, after controlling for age and gender. DMN state-specific positive BOLD responsiveness averaged across all seven DMN nodes was associated with the fit between the PDF of intervals between DMN electrophysiological activity states and the power law (negative of the deviation statistic in the upper panel; r=0.61, p<0.05). Follow-up analyses revealed that this result was driven by five nodes **(C)**; for all r-values, p<0.05; node numbers correspond to significant clusters in Figure 4. We found no association in control analyses between analogous BOLD responsiveness metrics for other HDR effects and corresponding interval-based power law statistics (abs(r)<0.38, p>0.16). Unlike the statistic computed based on DMN lifetimes, the power law fit estimated based on DMN intervals was not correlated with verbal or spatial episodic memory (rs<0.15 ps>0.61) but was correlated with global WMSA volumes **(D)**, which are an indicator of general cerebrovascular health. Differences in network connectivity patterns and cellular resource availability can both influence whether a network establishes critical behavior^106–108,195,196^. It is possible that these factors differentially modulate state lifetimes and intervals between coordinated activity states. Shaded areas, 95% confidence bounds.

